# Cross-continental variation of herbivore resistance in a global plant invader

**DOI:** 10.1101/2023.12.13.571471

**Authors:** Peipei Cao, Zhiyong Liao, Lei Zhang, Shengyu Wang, Jingwen Bi, Yujie Zhao, Madalin Parepa, Tiantian Lin, Yaolin Guo, Oliver Bossdorf, Christina L. Richards, Stacy B. Endriss, Jihua Wu, Ruiting Ju, Bo Li

## Abstract

- Successful plant invasions are often explained with adaptation to novel environments. However, invasive species often occupy broad niches within their native and introduced ranges, and a true understanding of microevolution during invasion therefore requires broad sampling of ranges, ideally with a knowledge of introduction history.
- We tested for genetic differentiation in herbivore resistance among 128 introduced (Europe, North America) and native (China, Japan) populations of the invasive Japanese knotweed (*Reynoutria japonica*) in two common gardens in the native range.
- In both common gardens we found that resistance traits of introduced populations differed from most Chinese native populations, but not from populations in Japan, the putative sources of introduction. Compared to Chinese populations, introduced European populations had thicker leaves with a lower C:N ratio but higher flavonoids contents. In the native range, variation in herbivore resistance was much more strongly associated with climate of origin than in introduced populations.
- Our results support the idea that founder effects played a key role in the invasion of knotweed into Europe and North America, with introduction of particular resistance phenotypes from Japan. Our study also demonstrates how knowledge of introduction history can avoid drawing wrong conclusions from observed biogeographic divergence.

## Introduction

The number of invasive plant species has increased dramatically over the past two centuries (Seebens *et al*., 2017), causing exceedingly negative impacts on environment (Powell *et al*., 2013; Bellard *et al*., 2016; Castro-Díez *et al*., 2019) and economy (Bradshaw *et al*., 2016; Diagne *et al*., 2021). In the context of globalization and climate change, one-sixth of the world’s land surface is expected to be highly vulnerable to invasion (Early *et al*., 2016), and the number of new invasive species is likely to increase further (Seebens *et al*., 2017, 2021) Thus, a comprehensive understanding of the mechanisms that underlie the successful invasion of alien species is an urgent issue in ecology and evolution (Kempel *et al*., 2013; Pyšek *et al*., 2020).

Many eco-evolutionary hypotheses proposed to explain increased performance of invasive populations are related to altered or novel abiotic or biotic environmental conditions in the introduced ranges (Cripps *et al*., 2006; Mitchell *et al*., 2006; Montti *et al*., 2016). The enemy release hypothesis (ERH) states that plant species experience a decrease in regulation by herbivores and other natural enemies, resulting in increased performance (Keane & Crawley, 2002). Since resistance to enemies can be costly (Koricheva, 2002), shifts in enemy pressure may lead to rapid evolutionary changes in growth and defense allocation (Bossdorf *et al*., 2005; Buswell *et al*., 2011; Li *et al*., 2022). The evolution of increased competitive ability (EICA) hypothesis therefore predicts that invasive populations may allocate fewer resources to herbivore defenses, and thus become more competitive than their native conspecifics (Blossey & Nötzold, 1995; Joshi & Vrieling, 2005; Lin *et al*., 2015). However, despite strong evidence for enemy release (Keane & Crawley, 2002; Liu & Stiling, 2006; Castells *et al*., 2013; Rotter *et al*., 2019; Xiao et al., 2020), especially with regard to specialists, the evidence in support of EICA is so far mixed (Bossdorf *et al*., 2005; Felker-Quinn *et al*., 2013; Rotter & Holeski, 2018), with lower herbivore resistance observed in several invasive species (Siemann & Rogers, 2001; Jakobs *et al*., 2004; Huang & Ding, 2016), but similar or even higher resistance in others (Genton *et al*., 2005; Lewis *et al*., 2006; Alba *et al*., 2011; Bhattarai *et al*., 2017; Gruntman *et al*., 2017; Lin *et al*., 2019). Some of these mixed results have been attributed to the different costs of different types of resistance (Koricheva, 2002; Neilson *et al*., 2013), and to the changes in herbivore community composition in the introduced ranges of invasive plants (Müller-Schärer *et al*., 2004; SDH-shifting defence hypothesis; Cripps *et al*., 2006).

Not only biotic conditions, but also altered abiotic environmental conditions may impose novel selection pressures and thus result in adaptive evolutionary changes in introduced populations (Buswell *et al*., 2011; Alexander *et al*., 2012; Lee & Kotanen, 2015). For instance, climatic conditions often differ between native and introduced ranges of invasive species (Early & Sax, 2014; Bocsi *et al*., 2016), and several previous studies have already demonstrated climate-related trait differentiation between native and introduced plant populations, suggesting rapid adaptation after introduction (Etterson *et al*., 2008; Alexander, 2013; Woods & Sultan, 2022). Furthermore, the existence of similar trait clines along climatic gradients in native and introduced populations is often also considered as evidence for adaptive post-invasion evolution (Hulme & Barrett, 2013; Bock *et al*., 2015), through multiple introductions of pre-adapted genotypes (Neuffer & Hurka, 1999; Keller & Taylor, 2008) and subsequent selection, i.e. ‘sorting out’ of standing genetic variation.

Climatic differences can also interact with the effects of herbivores. First, climate can directly affect plant defenses through physiological constraints and altered resource availability (Coley, 1985; Wright *et al*., 2004). Second, climatic conditions also influence the abundance and diversity of herbivores in plant communities (Maron *et al*., 2014; Anstett *et al*., 2016; Zhang *et al*., 2016a), thus indirectly driving selection for particular plant traits (Loughnan & Williams, 2019). Therefore, to understand invasive plant success and predict habitat vulnerability to future invasion, we need to understand the impacts of both enemy release and climatic conditions on herbivore resistance of invasive plant populations. Nevertheless, the relationships between climate and herbivore resistance have so far rarely been compared between native and introduced populations (Xiao *et al*., 2020).

Comparisons between native and introduced populations within common gardens are often facing several challenges (van Kleunen *et al*., 2018; Woods & Sultan, 2022). First, invasive species often occupy broad climatic niches both in their native and introduced ranges (Brandenburger *et al*., 2019, 2020; Liu *et al*., 2020), and comparisons of only few populations may lead to wrong conclusions, simply because the studied populations do not represent their respective ranges well enough (Yang *et al*.,2014). Second, when testing for evolution during invasion, the sources of introduced populations are of course the appropriate reference to invasive populations. Thus, a lack of knowledge of invasion history may further limit our ability to correctly identify trait changes in invasive populations (Cano *et al*., 2009; Bukovinszky *et al*., 2014). Third, because of widespread genotype-by-environment interactions (Richards *et al*., 2006), the results of common garden studies may to some extent be garden-specific. For instance, differences in abiotic and biotic conditions (e.g. climate, interacting species) might influence not only absolute values of growth and defense traits but also their relative differences between plant origins (Maron *et al*., 2004; Moloney *et al*., 2009; Qin *et al*., 2013; Yang *et al*., 2021).

Here, we worked with the invasive Japanese knotweed (*Reynoutria japonica*) and built on a previous cross-continental field survey of the species, which provided plant materials, along with important metadata, that allowed us to establish two large common gardens in different climatic zones of the native range, with 55 populations of origin from native range in China and Japan, and 73 populations from the introduced ranges in Europe and North America. The worldwide distribution and relatively clear invasion history of invasive knotweed provided an excellent model to explore biogeographic divergence in herbivore resistance and resistance traits (leaf traits and leaf chemistry), as well as their relationships with herbivore pressure and climatic factors at the collecting sites. We expected that (1) introduced populations have lower herbivore resistance than native populations, and that (2) traits of native populations are more strongly associated with climates of origin than those of introduced populations.

## Materials and methods

### Study system

*Reynoutria japonica* (Japanese knotweed) is native to eastern Asia and was introduced to Europe in the early 1840s and to North America in the 1870s as an ornamental (Bailey & Conolly, 2000). Along with its sister species *R. sachalinensis* it has become widely naturalized in both introduced ranges (Barney, 2006; Shimoda & Yamasaki, 2016; Del Tredici, 2017). *R. japonica*, *R. sachalinensis*, and their hybrid *R.* × *bohemica* spread rapidly along river banks and roadsides, often forming dense stands of hundreds of square meters (Bímová *et al*., 2004; Tiébré *et al*., 2008; Rouifed *et al*., 2014). Knotweed invasion seriously threatens the biodiversity and integrity of native ecosystems (Stoll *et al*., 2012; Mincheva *et al*., 2014), which causes substantial economic damage (Reinhardt *et al*., 2003; DEFRA, 2003). As a major environmental threat in Europe and North America, Japanese knotweed is listed among the 100 world’s worst invasive alien species by IUCN (Lowe *et al*., 2000).

### Sample collection

The plant materials we used came from a cross-latitudinal survey of 150 Japanese knotweeds populations in the native range of China and the introduced ranges of North America and Europe (Fig. **1**, Table S1; Irimia *et al*., 2023). In China our survey ranged from the Guangdong province in the South to the Shandong province in the (North-)East, in Europe from Northern Italy to Central Sweden, and in the United States from Georgia to Maine (Fig. **1**, Table S1). In each range we surveyed 50 populations along a 2000 km transect (*c.* every 40 km). The surveys in Europe and North America were done in 2019, that in China in 2020. For full details on the field survey please see Irimia *et al*. (2023). Briefly, at each sampling site we confirmed the knotweed taxon based on morphological characters (Bailey *et al*., 2009), laid a 30 m transect for sampling, selected five knotweed stems at regular intervals along the transect, and collected rhizomes from these individuals.

**Fig. 1.**
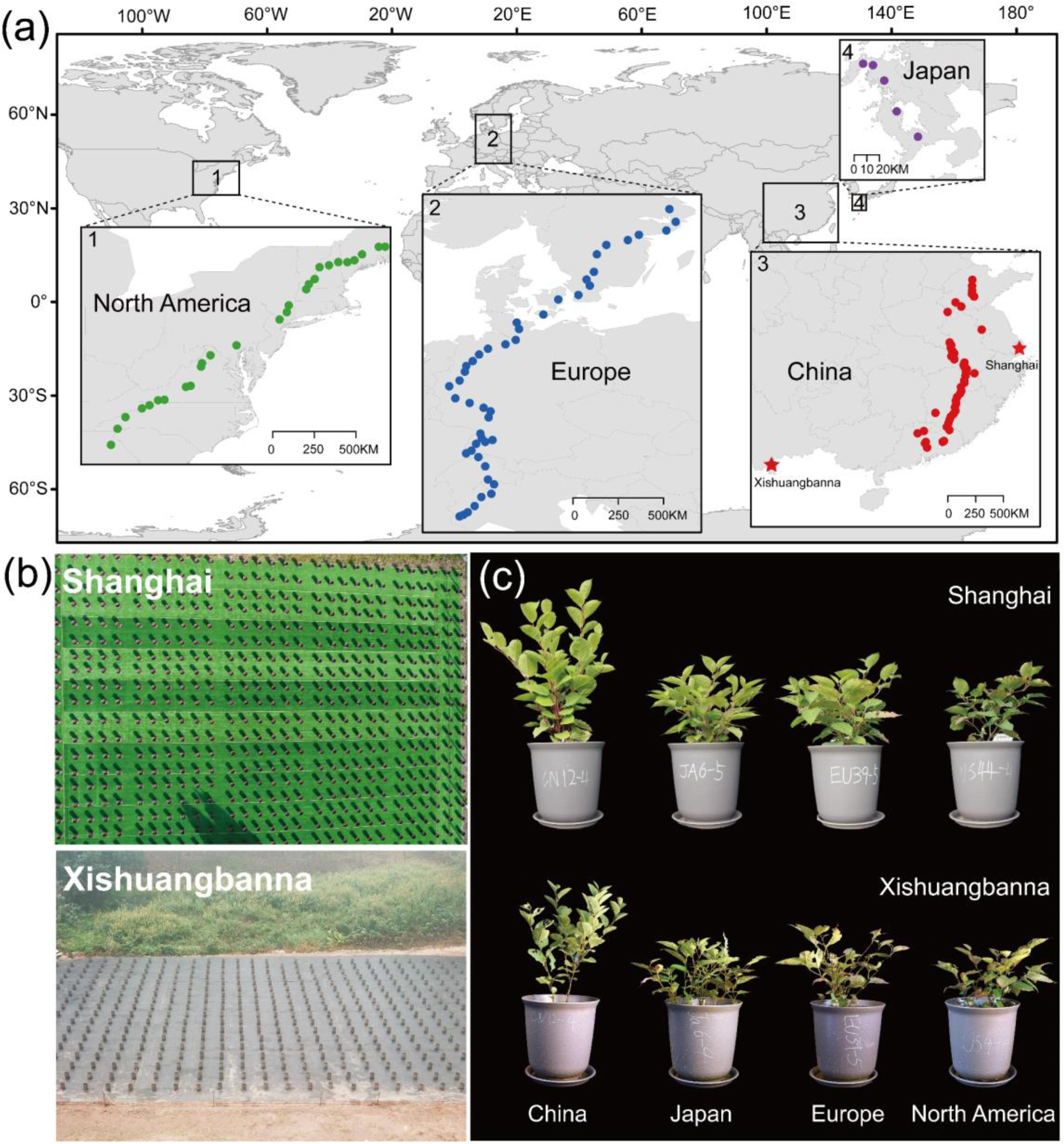
Geographic origins of the studied knotweed (*Reynoutria japonica*) clones, the two common gardens, and performance variation of different knotweed origins. (a) The geographic locations where rhizomes of *R. japonica* had been collected in the introduced ranges of North America (1; *n* = 27 populations) and Europe (2; *n* = 46), and in the native ranges of China (3; *n* = 50) and Japan (4; *n* = 5), and stars represent the locations of Shanghai (Fudan University Campus) and Xishuangbanna (Xishuangbanna Tropical Botanical Garden, Yunnan) common gardens. (b) Aerial photos of the two common garden experiments in Shanghai and Xishuangbanna. (c) Representative photos illustrating average growth differences between the two gardens and among differential ranges of origin in mid-July.

In addition to the main field survey, we also collected six native *R. japonica* populations from around Nagasaki in Japan, the region believed to be a key source of European and North American introductions (Bailey & Conolly, 2000; Del Tredici, 2017). A recent phylogenetic analysis based on chloroplast DNA (Zhang *et al*., unpublished data) has confirmed this hypothesis. Between 28 April and 4 May 2021, we collected rhizomes from 5-8 individuals separated by at least 6 m in each Japanese population. The rhizomes from Japan, Europe and North America were imported and temporally grown in a quarantine glasshouse at Xishuangbanna Tropical Botanical Garden, whereas the Chinese rhizomes were only stored at 4 °C for 2 weeks and then planted in pots to minimize maternal effects before setting up the common gardens. Because of legal restrictions, only one rhizome per North American populations could be imported prior to setting up the experiment.

### Common garden experiments

To test for heritable variation among knotweed plants from different ranges and populations, we set up two large common garden experiments, one at Fudan University in Shanghai (31°20′N, 121°30′E) and the other at Xishuangbanna Tropical Botanical Garden, Chinese Academy of Sciences (21°41′N, 101°25′E) in the Yunnan province. The two common garden locations differed in their climatic conditions, with warmer and moister conditions in Xishuangbanna (average March-October temperature 24.1 ℃; average monthly precipitation 171 mm) compared to Shanghai (average temperature 20.3 ℃; monthly precipitation 116 mm) (http://www.nmic.cn/; He *et al*., 2001; Song *et al*., 2010; Yu *et al*., 2021).

Because of import restrictions and variable cultivation success during quarantine conditions, we had to omit some introduced populations and ended up with a total of 128 *R. japonica* populations of origin: 55 populations from the native range (50 from China, 5 from Japan) and 73 populations from the introduced ranges (46 from Europe, 27 from North America) (Fig. **1**, Table S1). Both gardens were set up in open areas on flat ground covered with artificial grass mats.

Prior to planting the experiments, we cut rhizomes from each individual and kept them at 4 °C for at least four weeks to facilitate sprouting success. We cut all rhizomes to a size of 3-10 g (at least one intact node), removed the fine roots, and, as a measure of initial size, determined the fresh weight of each cutting. On 11 March 2022, all rhizomes were treated with fungicide and planted separately into 10.8 L pots filled with the same potting soil (Pindstrup substrate 0–10 mm, Pindstrup Mosebrug A/S, Denmark) in both gardens. Because of limited rhizome availability, we eventually planted a total of 518 *R. japonica* individuals in the Shanghai garden (27 individuals from 27 North Amercian populations; 218 individuals from 46 European populations; 239 individuals from 50 Chines populations; 34 individuals from 5 Japanese populations), and 463 individuals in the Xishuangbanna garden (16 individuals from 16 North American populations; 190 individuals from 46 European populations; 232 individuals from 50 Chinese populations; 25 individuals from 5 Japanese populations). In each garden, we arranged the plants in five blocks, ideally with one individual from each of the populations in each block, random assignment to blocks, and random order within blocks. To avoid aboveground interference, the distances between pots were all at least 90 cm. To avoid nutrient depletion, we added 10 g Osmocote fertilizer (Osmocote plus 801, N: P: K 16:8:12, Everris International B.V., Heerlen, Netherlands) to each pot at the beginning of the experiment, and once again in the middle of the experiment. Throughout the experiment, we watered the plants whenever the soil had become dry. To avoid losses of water and nutrients, all pots were individually placed on plastic trays. The pots, soils, and fertilizers used in the two common gardens were identical.

### Herbivore damage and resistance traits

To quantify variation in plant herbivore resistance, we estimated the degree of herbivore damage experienced by each individual in each garden, determined the levels of leaf secondary metabolites (lignin, flavonoids, alkaloids), and measured additional leaf traits that are often associated with palatability to herbivores (leaf toughness, leaf thickness, leaf ratios C : N) (Feng *et al*., 2011; Lin *et al*., 2015). In the middle of growing season (July 2022), we estimated herbivore damage as the percentage of leaf area eaten on each plant individual, separately for beetle and caterpillar damage (based on chewing modes and leaf notch types; Fig. S2). Two days later, we sampled five fresh, fully developed leaves (leaves 1, 2, 4, 5 and 6 from the top) from the tallest shoot of each plant individual, and measured the leaf thickness of each plant with a digital micrometer (Digimatic Outside Micrometer, Mitutoyo, Japan), and its toughness with a penetrometer (FA10, SAUTER, Balingen, Germany) in the Shanghai garden, and with a mechanical testing machine (ZQ990A, Dongguan Zhiqu Precision Instrument Co., Ltd, China) in the Xishuangbanna garden. For each plant, we then estimated the leaf thickness and toughness as the averages of the five measurements.

Finally, we dried all leaves at 60 °C for 72 h, and used these to analyze leaf chemistry. After grinding samples to the required particle size with a ball mill (MM400, Retsch, Germany), we measured total C and N with an organic elemental analyzer (FlashSmart™ Elemental Analyzer, Thermo-Fisher Scientific, USA) via thermal combustion and TCD/IR detection of CO^2^/N_2_, and also measured leaf lignin, alkaloids and total flavonoids using the MZS-1-G, SWJ-1-Y and LHT-1-G test kits (Suzhou Comin Biotechnology Co., Ltd., Suzhou, China), respectively. In total, we analysed 981 plant samples, 518 from the Shanghai garden and 463 individuals from the Xishuangbanna garden.

### Statistical analyses

To test for range differences in plant traits and herbivore damage, we fitted linear mixed models in R version 4.2.1 (R Core Team, 2022), with range (China, Japan, Europe, North America) as fixed effect, and population and block as random effects. To account for variation in initial size, our models included initial rhizome weight as covariates. We assessed the significance of fixed effects through Type Ⅲ Wald chi-squared tests using the *car* package (Fox & Weisberg, 2018). For traits that displayed a significant range effect (*P* < 0.05), we then conducted Tukey post-hoc tests with the *emmeans* and *pairs* functions (*emmeans* package; Lenth, 2018). Where necessary, we *log*-transformed the herbivore damage data to normalize the distribution of residuals.

To test for associations between leaf traits and herbivore damage, we calculated population-level spearman correlation coefficients for each pairwise combination of resistance trait (leaf traits and leaf chemistry) and herbivore damage, separately for each garden, using the *Hmisc* package (Hauke & Kossowski, 2011).

Finally, we tested whether the climatic conditions of population origins were associated with the resistance traits of native and invasive populations. For this, we obtained eight bioclimatic variables from the WorldClim database (Fick & Hijmans, 2017) that seemed particularly meaningful for characterising growing and overwintering conditions for knotweed plants: annual temperature (bio1), maximum temperature of the warmest month (bio4), minimum temperature of the coldest month (bio5), temperature seasonality (bio6), mean annual precipitation (bio12), precipitation of the wettest month (bio13), precipitation of the driest month (bio14), and precipitation seasonality (bio15). We simplified these climate data through a principal component analyses (PCA) using the *prcomp* function. The first and second principal component (‘climate PC1’ and ‘climate PC2’ hereafter) explained 54% and 21% of the variance in the eight climatic variables across all populations, respectively, with climate PC1 mainly related to mean temperature as well as cold and precipitation extremes, and climate PC2 related to temperature maxima and seasonality (Fig. S1, Table S2). We then performed population-level linear regression analyses with either climate PC1 or climate PC2 as explanatory variables, and the common-garden averages of the resistance traits of native or introduced populations as dependent variables (Moreira *et al*., 2018; Galmán *et al*., 2021).

## Results

We found that on average the traits of invasive populations often differed from those of native populations. However, these overall patterns were largely driven by the native populations from China which were distinct from European and North American populations, whereas the native populations from Japan were often similar to the invasive ones.

### Range differences in herbivore damage

The average levels of herbivore damage differed substantially between the two common gardens: in Xishuangbanna the percentages of leaf area eaten by beetles and caterpillars were three and 19 times higher, respectively, than in Shanghai (Fig. **2**). In Shanghai, plants from Chinese populations experienced significantly higher levels of caterpillar herbivory than plants from Europe or North America (Fig. **2b**), and the same was true for beetle damage in Xishuangbanna, where the Chinese populations were also most strongly attacked (Fig. **2c**). Interestingly, Chinese populations experienced less caterpillar damage than European populations in Xishuangbanna, but we never found any significant differences between the two introduced ranges, or between Japanese populations and the invasive populations, in the two gardens.

**Fig. 2.**
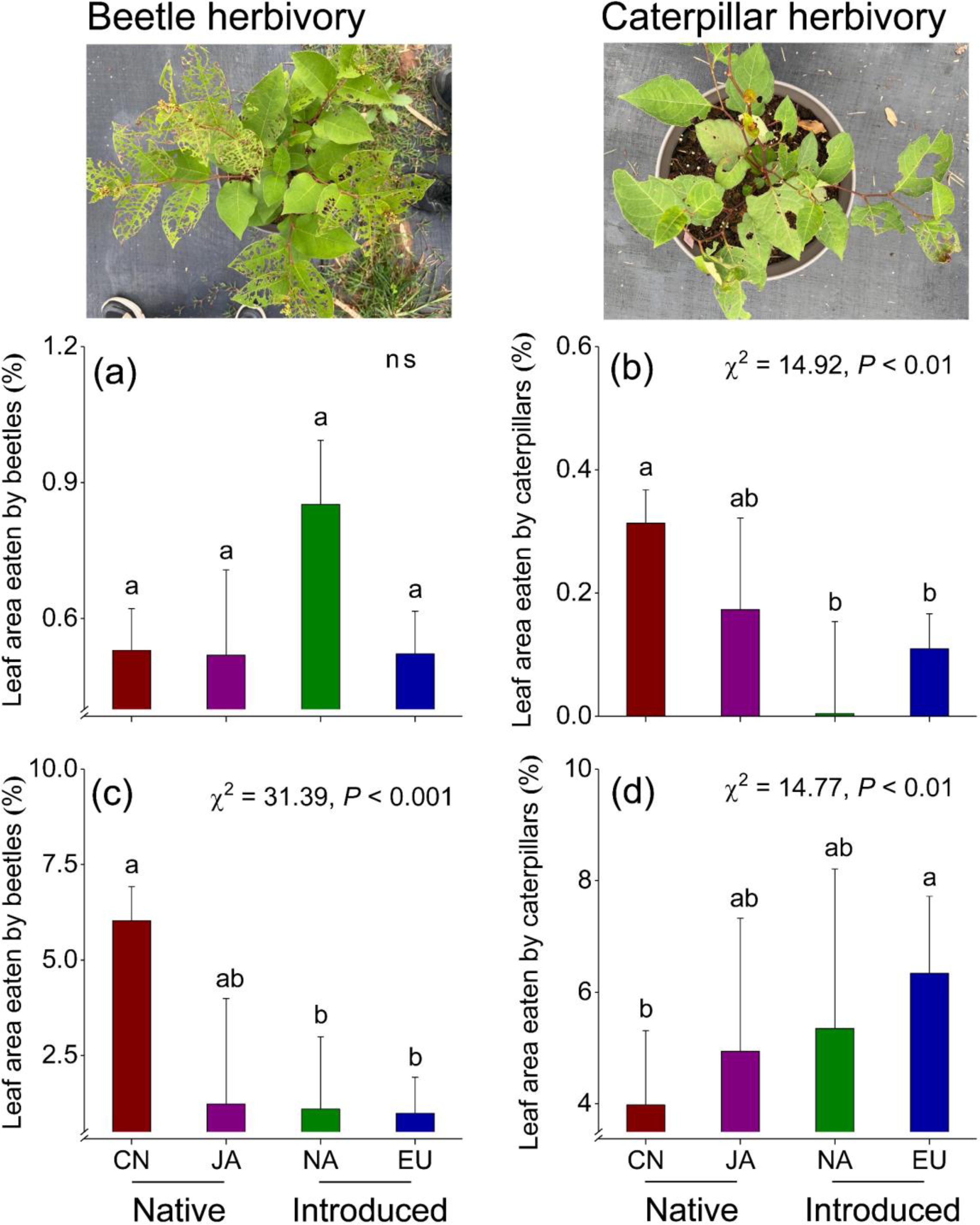
Average levels of herbivores damage observed in *R. japonica* plants from China (CN), Japan (JA), North America (NA) and Europe (EU) when grown in common gardens in Shanghai (a, b) and Xishuangbanna (c, d). The values are adjusted means and SEs from ANOVA, with different letters above error bars indicating significant group differences based on Tukey’s HSD post-hoc tests. *Chi-square* statistics and significance level of regression models are also shown. ns, not significant.

### Range differences in resistance traits

Plants from native Chinese populations were also distinct in their resistance traits. In both common gardens, Chinese plants had significantly thinner leaves than plants from invasive populations (Fig. **3a,d**), and in the Shanghai garden they also had higher leaf ratios C : N and lower leaf flavonoids than plants from both invasive ranges (Figs. **3c** and **4c**). Moreover, Chinese populations had tougher leaves than European populations in the Shanghai garden (Fig. **3b**) and higher leaf ratios C : N than North American populations in the Xishuangbanna garden (Fig. **3f**). We did not find any range differences in leaf lignin and leaf alkaloids between native and invasive populations Fig. **4a,b****,d,e**), and there were no significant differences in leaf traits between native populations from Japan and the invasive populations (Figs. **3** and **4**).

**Fig. 3.**
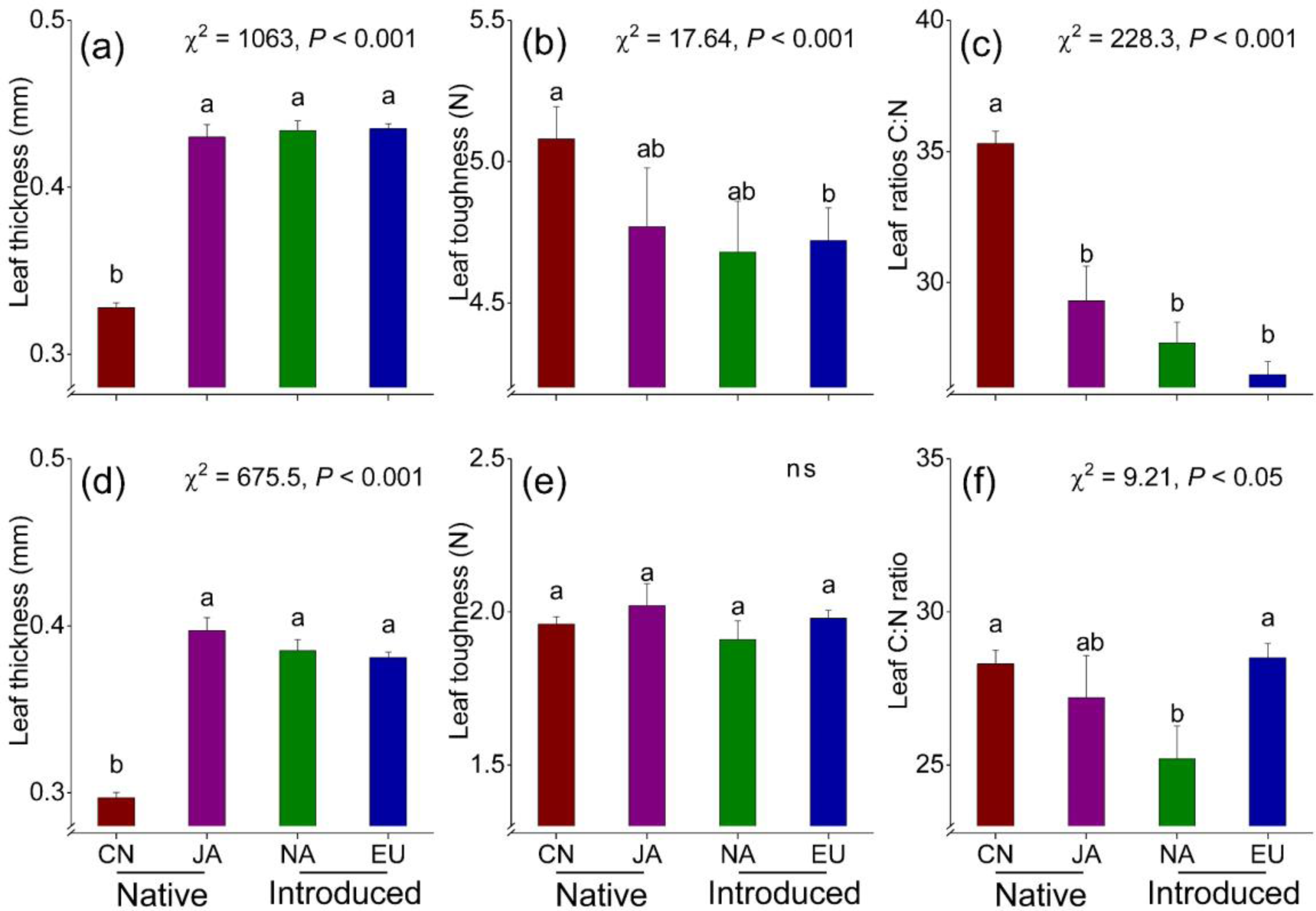
Average levels of leaf traits observed in *R. japonica* plants from China (CN), Japan (JA), North America (NA) and Europe (EU) when grown in common gardens in Shanghai (a-c) and Xishuangbanna (d-f). The values are adjusted means and SEs from ANOVAs, with different letters above error bars indicating significant group differences based on Tukey’s HSD post-hoc tests. *Chi-square* statistics and significance level of regression models are also shown. ns, not significant.

**Fig. 4.**
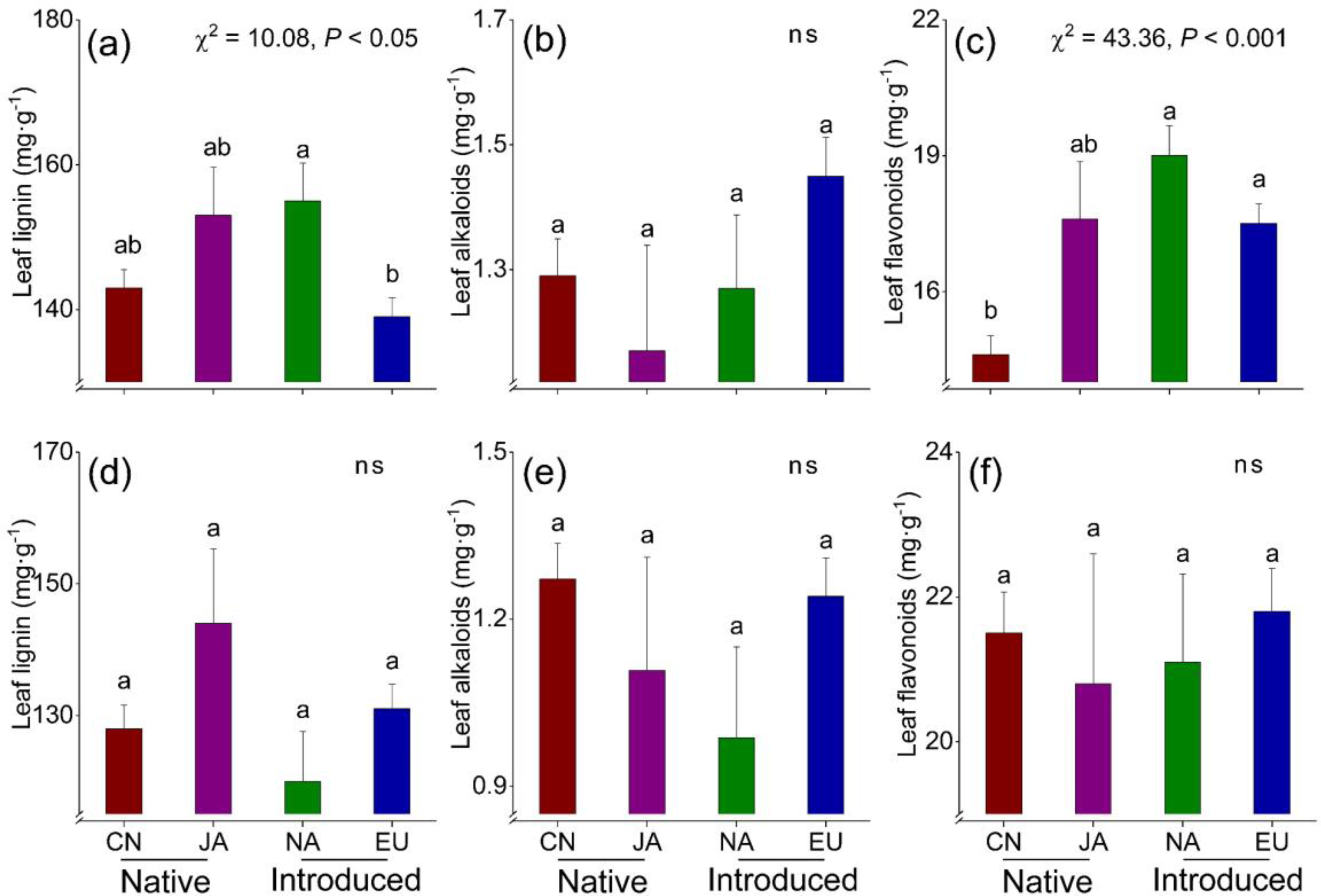
Average levels of leaf chemistry observed in *R. japonica* plants from China (CN), Japan (JA), North America (NA) and Europe (EU) when grown in common gardens in Shanghai (a-c) and Xishuangbanna (d-f). The values are adjusted means and SEs from ANOVAs, with different letters anove error bars indicate significant group differences based on Tukey’s HSD post-hoc tests. *Chi-square* statistics and significance level of regression models are also shown. ns, not significant.

### Correlations between resistance traits and herbivore damage

In the Shanghai garden, leaf damage by caterpillars was negatively correlated with leaf thickness and leaf flavonoids, but positively with leaf ratios C : N (Table 1). In contrast, in the Xishuangbanna garden, beetle herbivory was negatively correlated with leaf thickness and flavonoids, but caterpillar herbivory was positively correlated with leaf thickness (Table 1).

**Table 1.** Spearman’s rank correlations (*r_s_*) between leaf traits and leaf herbivory, tested across 128 native and introduced populations of Japanese knotweed, when grown in two different common gardens in Shanghai and Xishuangbanna

|  | Thickness | Toughness | Ratios C : N | Lignin | Alkaloids | Flavonoids |
| --- | --- | --- | --- | --- | --- | --- |
| *Shanghai* |  |  |  |  |  |  |
| % Leaf area eaten by beetles | 0.13 | -0.06 | -0.11 | -0.11 | 0.10 | -0.07 |
| % Leaf area eaten by caterpillars | **-0.40\*\*\*** | 0.12 | **0.34\*\*\*** | -0.14 | 0.05 | **-0.35\*\*\*** |
| *Xishuangbanna* |  |  |  |  |  |  |
| % Leaf area eaten by beetles | **-0.30\*\*** | -0.13 | -0.06 | 0.06 | 0.00 | **-0.31\*\*\*** |
| % Leaf area eaten by caterpillars | **0.19\*** | -0.08 | -0.03 | 0.15 | 0.03 | 0.11 |

Significant correlations are in bold. Significance levels: \*, $P < 0.05$ ; \*\*, $P < 0.01$ ; \*\*\*, $P < 0.001$ .

### Correlations between climates of origin and resistance traits

Significant correlations between climates of origin and resistance traits were frequent among native populations but much less common among introduced populations (10 vs. 3 correlations in native vs introduced plants; Table 2). In both common gardens, leaf thickness was negatively correlated with climate PC1 but positively correlated with climate PC2 for native but not invasive plants, and leaf ratios C : N of native populations was negatively related to climate PC1. Only in the Xishuangbanna garden, leaf ratios C : N of invasive populations was positively correlated to climate PC2. For native populations, leaf flavonoids were positively related to climate PC1 but negatively to PC2 in both gardens, but for invasive populations we found a positive climate PC1-flavonoid correlation only in Xishuangbanna, and a negative correlation of flavonoids with climate PC2 only in Shanghai (Table 2).

**Table 2.**
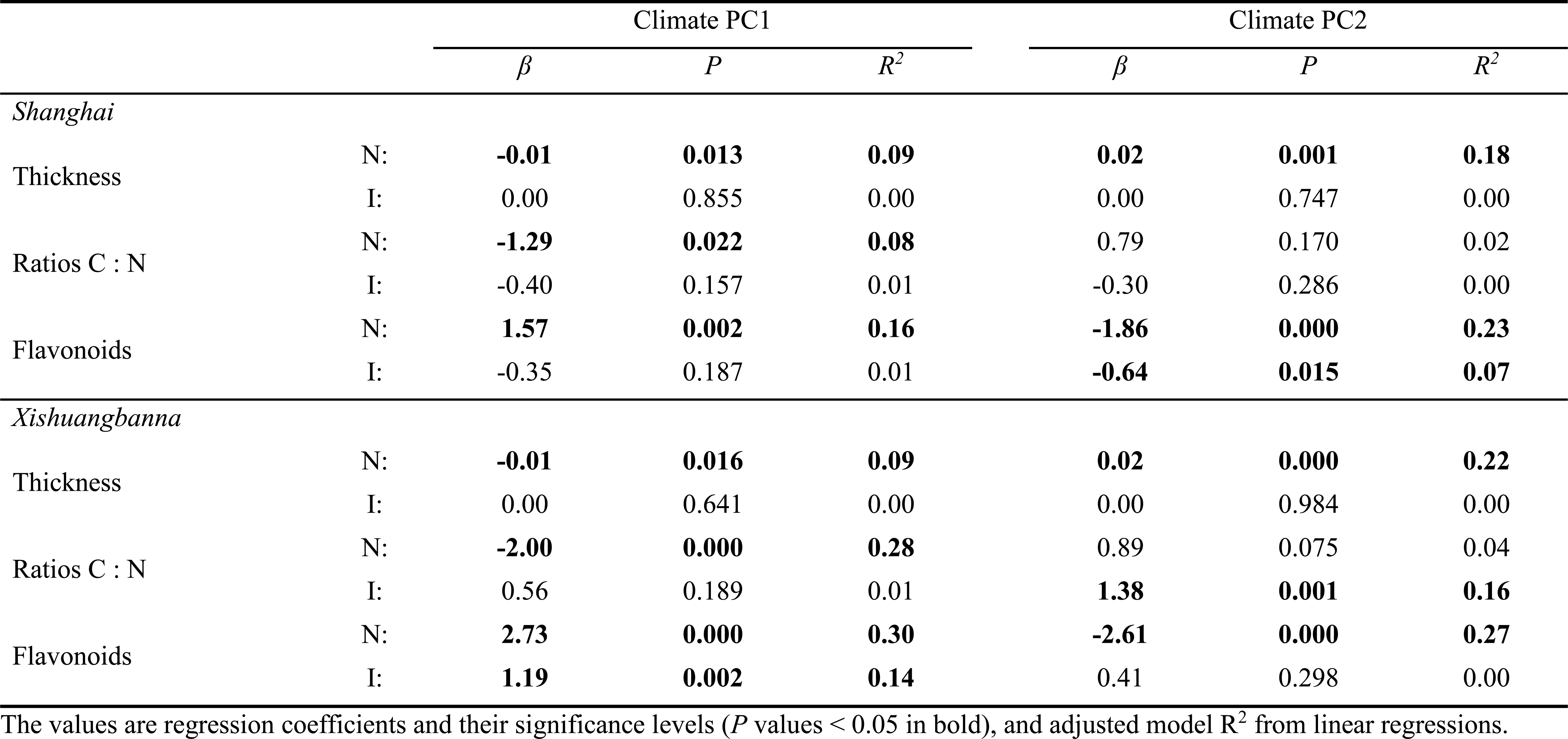
Associations between climates of origin (first two PCs from a PCA of eight bioclimatic variables, see Table S2) and variation in leaf traits across native (N; China and Japan) and introduced (I; Europe and North America) Japanese knotweed populations, when grown in two common gardens in Shanghai and Xishuangbanna in China

In other words, native populations from warmer, humid sites with high temperature seasonality tended to have thicker leaves with a higher ratios C : N and lower flavonoids contents (Table 2), where for invasive populations climate associations were rarer and more variable: in Shanghai, higher flavonoid contents of invasive populations were associated with high summer temperature and low seasonality whereas in Xishuangbanna lower flavonoids were associated with warm and humid areas of origin (Table 2).

## Discussion

Successful plant invasions are often explained by evolutionary changes in plant traits. However, the power of previous common garden comparisons of native versus introduced populations has often remained limited. Here, we tested for genetic differentiation in herbivore resistance among 128 native and introduced populations of invasive knotweed (*Reynoutria japonica*) in two common gardens in the native range. We found that on average introduced populations from Europe and North America differed in several resistance traits from native Asian populations, but that these overall differences were driven by native populations from China, whereas the resistance traits of native Japanese populations were generally similar to populations from the introduced range. Our study supports – at the level of phenotypes – a Japanese source of knotweed invasions, and that founder effects – rather than post-invasion evolution – explains overall range differences in knotweed traits.

### Range differences in herbivore resistance

The EICA hypothesis predicts that because of a release from native herbivores introduced populations might have reduced herbivore defenses (Blossey & Notzold, 1995). We found that introduced European and North American populations were indeed significantly different in many leaf traits from native Chinese populations, but that there were little differences between introduced populations and the putative source populations from Japan. Our results thus demonstrate the importance of accounting for invasion history when testing for evolution during invasions (Colautti & Lau, 2015; Brandenburger *et al*., 2020).

We found no significant differences in many herbivore resistance traits such as leaf ratios C : N, leaf toughness, leaf flavonoids, lignin or alkaloids between introduced European and North American and native Japanese populations in both common gardens. There were also no differences in the herbivore damage of plants in both gardens, which represented two different natural herbivore environments in the native range. Our results thus did not support the EICA prediction that invasive populations have evolved lower herbivore resistance than their native sources (Blossey & Notzold, 1995). Several studies with other invasive species have shown no difference or even higher resistance of invasive populations in common garden experiments, e.g. with *Chromolaena odorata* (Liao *et al*., 2014), *Brassica nigra* (Oduor *et al*., 2011), or *Verbascum thapsus* (Endriss *et al*., 2018). We are only aware of one previous common garden study with native and introduced populations of *R. japonica* (Rouifed *et al*. 2018). It compared only five native Japanese and five invasive French populations in a common garden in the introduced range, but it also found that introduced populations did not differ in their composition of secondary metabolites, stem stiffness and leaf thickness, but that they had even higher leaf toughness than native Japanese populations. As the Japanese populations used in our experiments – as well as those used in Rouifed *et al*. (2018) – were from the putative source region of knotweed introductions to Europe and North America, our results thus indicate that the introduction of plants with particular resistance profiles, rather than post-invasion evolution, underlies the trait divergence between the native and introduced range, and may have played a role in the invasion success of *R. japonica*.

We found that herbivore damage varied substantially between the two common gardens, and related to different leaf traits. For instance, in the Shanghai garden damage by caterpillars was negatively associated with leaf thickness and flavonoids, but positively with leaf ratios C : N, whereas in Xishuangbanna, the same type of damage was only positively associated with leaf thickness. Part of these divergent results might have resulted from the different herbivore communities in the two common gardens, e.g. if different herbivores have different feeding preferences (Van der Meijden, 1996; Bossdorf *et al*., 2004; Yang *et al*., 2014). In Xishuangbanna, we found that damage by beetles was associated with different traits than damage by caterpillars: beetles preferred plants with thinner leaves and lower flavonoid levels, whereas caterpillars preferred plants with thicker leaves. Besides the herbivore communities, the two common gardens of course also differed in climatic conditions, including temperature, precipitation and solar radiation, and since leaf traits could also be related to other environmental drivers, e.g. influence drought or UV tolerance (Levitt & Lovett, 1985; Strauss & Agrawal, 1999; Harborne & Williams, 2000; Peter Constabel *et al*., 2014; Barton & Boege, 2017; Li *et al*., 2022), this may have further contributed to divergent herbivory-leaf trait relationships. More generally, our results demonstrate the environmental contingency of common garden results, including when testing for variation in herbivore resistance (Maron *et al*., 2004; Qin *et al*., 2013; Yang *et al*., 2014; Bossdorf *et al*., 2005; Yang *et al*., 2021), and hence the value of working with multiple common gardens.

### Associations between climates of origin and herbivore resistance

We found that the climatic conditions at the collecting sites were significantly associated with resistance traits in both common gardens, but only for plants from the native range, whereas for plants from the introduced range there were only very few significant associations. Many previous studies of invasive plants have documented the development of parallel clinal patterns in ecological traits among populations of native and introduced ranges (Agrawal *et al*., 2015; Leger & Rice, 2007; Etterson *et al*., 2008; Rosche *et al*., 2019; Hodgins *et al*., 2020) and have usually interpreted this as evidence for rapid post-invasion evolution and adaptation. However, there are also studies of successful invasive plants with unparalleled (Bhattarai *et al*., 2017; Yang *et al*., 2021; Woods & Sultan, 2022) or no clinal patterns at all (Endriss *et al*., 2018; Sun & Roderick, 2019), and our results also provide little evidence for post-invasion genetic differentiation of introduced European or North American populations in relation to climate.

A simple reason for the observed lack of genetic differentiation could be that there was too little genetic variation for natural selection to act on. The majority of invasive *R. japonica* populations are thought to be descendants of a single introduced clone (Hollingsworth & Bailey, 2000; Richards *et al*., 2012; Gaskin *et al*., 2014; Groeneveld *et al*., 2014; Zhang *et al*., 2016b). However, so far this knowledge is based on low-resolution molecular studies of limited numbers of populations, and we clearly need broader and higher-resolution population genomic analyses to evaluate this explanation. Moreover, even if little genetic variation was introduced in the first place, novel mutations can sometimes be common enough to result in genetic variation that can be exploited for adaptation, even within a relatively short period of time after an invasion (Lynch & Conery, 2000, Ossowski *et al*., 2010). In any case, an alternative explanation for the lack of climate-related genetic differentiation could be that adaptation did not occur *yet*. Although rapid evolutionary changes have frequently been reported for invasive plants, examples from the invasion literature show that geographic clines usually develop 50-150 generations after introduction (Moran & Alexander, 2014), but many of these examples came from short-lived plants. *Reynoutria japonica* was introduced to Europe *c.* 170 years ago and some decades later to North America, but it began to expand rapidly only in the past 50-100 years (Bailey & Conolly, 2000), and it is a perennial, so it is a possibility that there was not enough time. Again, genomic approaches will help to answer this question.

## Conclusion

Our study combined replicate common gardens and a biogeographical approach with samples spanning an exceptionally large climatic gradient from both introduced and native ranges to understand evolution of herbivore resistance in the invasive species *R. japonica*. We found that the resistance traits of introduced European and North American populations differed from most Chinese native populations but were generally similar to Japanese populations that are putative sources of the introductions. Thus, we find little evidence for post-invasion evolution but that plants with particular resistance profiles have been introduced to Europe and North America, and may have played a role in the invasion success of *R. japonica*. Our study demonstrates the importance of accounting for invasion history when testing for evolution during invasion.

## Acknowledgements

We thank Fatima Elkott, Christiane Karasch-Wittmann, Elodie Kugler, Rongjin Li, Kaitao Liu, Rui Min, Jeannie Mounger, Julia Rafalski, Conner Richardson, Eva Schloter, Sabine Silberhorn, Weihan Zhao, Wenchao Zhong, Zhiming Zhong and Xin Zhuang for help with field collections, and Jieren Jin, Guangye Li, Shibo Tian, Chaonan Wang and Niyao Xiang with common garden experiments. We also thank the Center for Gardening and Horticulture and the Seed Bank of Tropical Plant Resources at XTBG for providing quarantine facilities and cold rooms. This study was supported by the National Natural Science Foundation of China (grant 31961133028), the German Federal Ministry of Education and Research (MOPGA Project 306055) and the German Research Foundation (grant 431595342).

## Author contributions

PC, ZL, MP, OB, CLR and BL designed the experiments. PC, ZL, SW, LZ, YZ and JB conducted the research. PC and ZL analyzed the data and wrote the first draft. All authors contributed to the revision and improvement of the manuscript.

## Data availability statement

Data available from the Dryad Digital Repository (private for peer review): https://datadryad.org/stash/share/EYcQ6ThtOzgWS6oC2dS3Xlj7ZhylK9veRFDySOd EZQ0, DOI is 10.5061/dryad.qbzkh18r4.

## Supporting Information

**Table S1.** Geographic origins of the 128 studied *Reynoutria japonica* populations and their uses in the Shanghai (SH) and Xishuangbanna (XSBN) common gardens.

| Population ID | Site | Latitude | Longitude | Garden |
| --- | --- | --- | --- | --- |
| *China (Native)* |  |  |  |  |
| CN01 | Guangdong | 23.29 | 114.01 | SH / XSBN |
| CN02 | Guangdong | 23.64 | 113.84 | SH / XSBN |
| CN03 | Guangdong | 23.74 | 113.91 | SH / XSBN |
| CN04 | Guangdong | 23.75 | 115.27 | SH / XSBN |
| CN05 | Guangdong | 23.82 | 115.38 | SH / XSBN |
| CN06 | Guangdong | 24.44 | 113.25 | SH / XSBN |
| CN07 | Guangdong | 24.62 | 113.76 | SH / XSBN |
| CN08 | Guangdong | 24.71 | 115.82 | SH / XSBN |
| CN09 | Jiangxi | 24.96 | 115.61 | SH / XSBN |
| CN10 | Jiangxi | 25.24 | 115.73 | SH / XSBN |
| CN11 | Jiangxi | 25.53 | 115.84 | SH / XSBN |
| CN12 | Jiangxi | 25.80 | 115.94 | SH / XSBN |
| CN13 | Jiangxi | 26.03 | 116.16 | SH / XSBN |
| CN14 | Jiangxi | 26.10 | 114.69 | SH / XSBN |
| CN15 | Jiangxi | 26.27 | 116.32 | SH / XSBN |
| CN16 | Jiangxi | 26.55 | 116.27 | SH / XSBN |
| CN17 | Jiangxi | 26.85 | 116.37 | SH / XSBN |
| CN18 | Jiangxi | 27.08 | 116.34 | SH / XSBN |
| CN19 | Jiangxi | 27.38 | 116.45 | SH / XSBN |
| CN20 | Jiangxi | 27.66 | 116.68 | SH / XSBN |
| CN21 | Jiangxi | 27.87 | 116.78 | SH / XSBN |
| CN22 | Jiangxi | 28.19 | 116.76 | SH / XSBN |
| CN23 | Jiangxi | 28.54 | 117.05 | SH / XSBN |
| CN24 | Jiangxi | 28.71 | 117.02 | SH / XSBN |
| CN25 | Jiangxi | 29.10 | 117.12 | SH / XSBN |
| CN26 | Jiangxi | 29.21 | 117.15 | SH / XSBN |
| CN27 | Jiangxi | 29.31 | 117.85 | SH / XSBN |
| CN28 | Jiangxi | 29.43 | 117.16 | SH / XSBN |
| CN29 | Jiangxi | 29.64 | 117.22 | SH / XSBN |
| CN30 | Anhui | 29.81 | 117.05 | SH / XSBN |
| CN31 | Anhui | 29.98 | 116.97 | SH / XSBN |
| CN32 | Anhui | 30.18 | 117.04 | SH / XSBN |
| CN33 | Anhui | 30.44 | 116.23 | SH / XSBN |
| CN34 | Anhui | 30.69 | 115.98 | SH / XSBN |
| CN35 | Anhui | 30.82 | 116.05 | SH / XSBN |
| CN36 | Anhui | 30.92 | 116.21 | SH / XSBN |
| CN37 | Anhui | 31.09 | 116.10 | SH / XSBN |
| CN38 | Anhui | 31.21 | 116.02 | SH / XSBN |
| CN39 | Anhui | 31.40 | 115.93 | SH / XSBN |
| CN40 | Anhui | 31.58 | 115.97 | SH / XSBN |
| CN41 | Henan | 31.81 | 115.85 | SH / XSBN |
| CN42 | Jiangsu | 32.85 | 118.44 | SH / XSBN |
| CN43 | Henan | 34.27 | 115.68 | SH / XSBN |
| CN44 | Jiangsu | 34.72 | 116.79 | SH / XSBN |
| CN45 | Shandong | 35.06 | 116.31 | SH / XSBN |
| CN46 | Shandong | 35.52 | 117.82 | SH / XSBN |
| CN47 | Shandong | 35.74 | 117.64 | SH / XSBN |
| CN48 | Shandong | 36.00 | 117.65 | SH / XSBN |
| CN49 | Shandong | 36.38 | 117.66 | SH / XSBN |
| CN50 | Shandong | 36.87 | 117.68 | SH / XSBN |
| *Japan (Native)* |  |  |  |  |
| JA01 | Unzen | 32.81 | 129.92 | SH / XSBN |
| JA02 | Saikai | 32.99 | 129.77 | SH / XSBN |
| JA03 | Sasebo | 33.21 | 129.68 | SH / XSBN |
| JA04 | Hirado | 33.32 | 129.60 | SH / XSBN |
| JA05 | Hirado | 33.33 | 129.53 | SH / XSBN |
| *North America (Introduced)* |  |  |  |  |
| NA01 | Georgia | 34.24 | -83.46 | SH / XSBN |
| NA02 | NorthCarolina | 35.10 | -83.10 | SH / XSBN |
| NA03 | NorthCarolina | 35.74 | -82.68 | SH / XSBN |
| NA04 | NorthCarolina | 36.21 | -81.78 | SH / XSBN |
| NA05 | NorthCarolina | 36.38 | -81.38 | SH / XSBN |
| NA06 | Virginia | 36.65 | -80.92 | SH / XSBN |
| NA07 | Virginia | 36.67 | -80.57 | SH |
| NA08 | Virginia | 37.37 | -79.41 | SH |
| NA09 | Virginia | 37.43 | -79.16 | SH |
| NA10 | Virginia | 38.46 | -78.59 | SH / XSBN |
| NA11 | Virginia | 38.65 | -78.54 | SH |
| NA12 | Virginia | 39.08 | -78.09 | SH / XSBN |
| NA13 | Maryland | 39.61 | -76.68 | SH / XSBN |
| NA14 | NewJersey | 41.01 | -74.35 | SH / XSBN |
| NA15 | NewYork | 41.41 | -73.96 | SH / XSBN |
| NA16 | NewYork | 41.77 | -73.86 | SH |
| NA17 | Massachusetts | 42.64 | -72.91 | SH / XSBN |
| NA18 | Vermont | 42.92 | -72.77 | SH / XSBN |
| NA19 | Vermont | 43.20 | -72.45 | SH / XSBN |
| NA20 | Vermont | 43.84 | -72.19 | SH |
| NA21 | NewHampshire | 43.94 | -71.68 | SH / XSBN |
| NA22 | NewHampshire | 44.12 | -71.18 | SH / XSBN |
| NA23 | Maine | 44.11 | -70.69 | SH |
| NA24 | Maine | 44.21 | -70.31 | SH |
| NA25 | Maine | 44.53 | -69.89 | SH |
| NA26 | Maine | 44.94 | -68.99 | SH |
| NA27 | Maine | 44.95 | -68.64 | SH |
| *Europe (Introduced)* |  |  |  |  |
| EU01 | Italy | 44.67 | 7.29 | SH / XSBN |
| EU02 | Italy | 44.75 | 7.48 | SH / XSBN |
| EU03 | Italy | 44.88 | 7.69 | SH / XSBN |
| EU04 | Italy | 45.19 | 8.04 | SH / XSBN |
| EU05 | Italy | 45.64 | 8.37 | SH / XSBN |
| EU06 | Italy | 45.80 | 8.87 | SH / XSBN |
| EU07 | Switzerland | 46.27 | 9.00 | SH / XSBN |
| EU08 | Switzerland | 46.51 | 8.7 | SH / XSBN |
| EU09 | Switzerland | 47.16 | 8.56 | SH / XSBN |
| EU10 | Germany | 47.62 | 8.22 | SH / XSBN |
| EU11 | Germany | 47.81 | 7.61 | SH / XSBN |
| EU12 | Germany | 47.94 | 7.88 | SH / XSBN |
| EU13 | Germany | 48.28 | 8.11 | SH / XSBN |
| EU14 | Germany | 48.37 | 8.56 | SH / XSBN |
| EU15 | Germany | 48.47 | 8.92 | SH / XSBN |
| EU16 | Germany | 48.56 | 8.39 | SH / XSBN |
| EU17 | Germany | 48.79 | 8.32 | SH / XSBN |
| EU18 | Germany | 49.59 | 8.73 | SH / XSBN |
| EU19 | Germany | 49.90 | 8.83 | SH / XSBN |
| EU20 | Germany | 50.07 | 8.48 | SH / XSBN |
| EU21 | Germany | 50.31 | 7.79 | SH / XSBN |
| EU22 | Germany | 50.54 | 7.08 | SH / XSBN |
| EU23 | Germany | 51.14 | 6.78 | SH / XSBN |
| EU24 | Germany | 51.43 | 7.29 | SH / XSBN |
| EU25 | Germany | 51.87 | 7.55 | SH / XSBN |
| EU26 | Germany | 52.15 | 7.62 | SH / XSBN |
| EU27 | Germany | 52.39 | 7.94 | SH / XSBN |
| EU28 | Germany | 52.72 | 8.26 | SH / XSBN |
| EU29 | Germany | 53.01 | 8.70 | SH / XSBN |
| EU30 | Germany | 53.22 | 9.57 | SH / XSBN |
| EU31 | Germany | 53.45 | 10.08 | SH / XSBN |
| EU32 | Germany | 53.99 | 10.25 | SH / XSBN |
| EU33 | Germany | 54.31 | 10.13 | SH / XSBN |
| EU34 | Denmark | 54.71 | 11.45 | SH / XSBN |
| EU35 | Denmark | 55.46 | 12.19 | SH / XSBN |
| EU36 | Sweden | 55.68 | 13.19 | SH / XSBN |
| EU37 | Sweden | 56.14 | 13.76 | SH / XSBN |
| EU38 | Sweden | 56.45 | 13.60 | SH / XSBN |
| EU39 | Sweden | 56.83 | 13.96 | SH / XSBN |
| EU40 | Sweden | 57.70 | 14.11 | SH / XSBN |
| EU41 | Sweden | 58.17 | 14.58 | SH / XSBN |
| EU42 | Sweden | 58.41 | 15.65 | SH / XSBN |
| EU43 | Sweden | 58.67 | 16.20 | SH / XSBN |
| EU44 | Sweden | 58.89 | 17.56 | SH / XSBN |
| EU45 | Sweden | 59.32 | 18.02 | SH / XSBN |
| EU46 | Sweden | 59.95 | 17.71 | SH / XSBN |

**Table S2.**
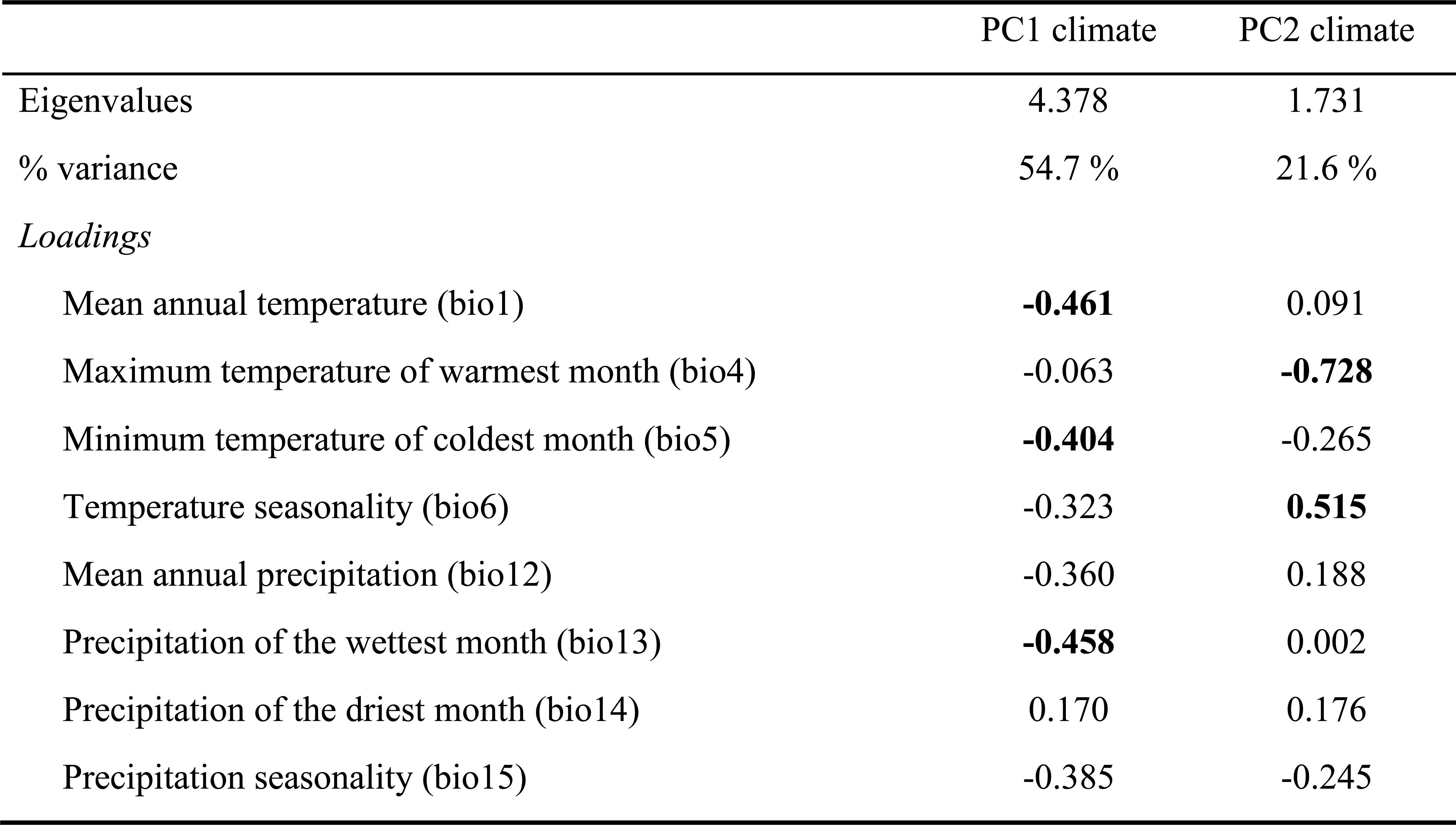
Summary of PCA of eight climate variables. The first and second components explained 54% and 21% of climatic variation among native and introduced ranges’ localities, respectively. Climate variables with stronger loadings (> 0.40) on climate PCs are shown in bold.

**Fig. S1.**
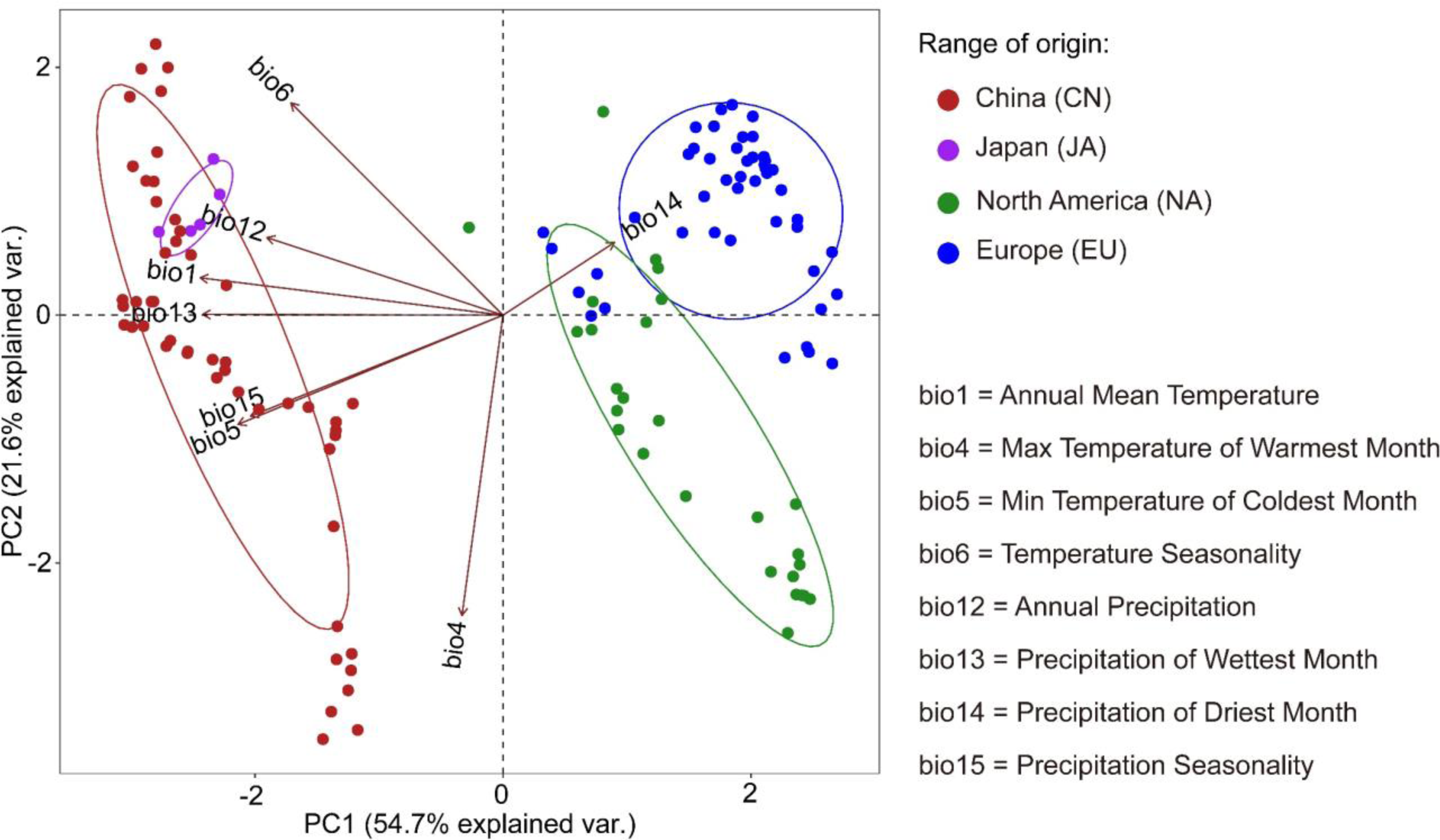
Principal component analysis (PCA) for climate variables at sample collecting sites of *R. japonica* in common gardens. Colored dots indicate sample collecting sites. Percentages on the X- and Y-axes indicate the variation explained by each principal component.

